# pr-independent biogenesis of infectious mature Zika virus particles

**DOI:** 10.1101/2024.09.12.612520

**Authors:** Kimberly A. Dowd, Michelle Schroeder, Egan Sanchez, Beniah Brumbaugh, Bryant M. Foreman, Katherine E. Burgomaster, Wei Shi, Lingshu Wang, Natalie Caputo, David N. Gordon, Cindi L. Schwartz, Bryan T. Hansen, Maya Aleshnick, Wing-Pui Kong, Kaitlyn M. Morabito, Heather D. Hickman, Barney S. Graham, Elizabeth R. Fischer, Theodore C. Pierson

## Abstract

Flavivirus assembly at the endoplasmic reticulum is driven by the structural proteins envelope (E) and premembrane (prM). Here, contrary to the established paradigm for flavivirus assembly, we demonstrate that the biogenesis of flavivirus particles does not require an intact prM nor proteolytic activation. The expression of E preceded by a truncated version of prM (M-E) was sufficient for the formation of non-infectious Zika virus subviral particles and pseudo-infectious reporter virions. Subviral particles encoded by a ZIKV M-E DNA vaccine elicited a neutralizing antibody response that was insensitive to the virion maturation state, a feature of flavivirus humoral immunity shown to correlate with protection. M-E vaccines that uniformly present structural features shared with mature virions offer a higher quality and broadly applicable approach to flavivirus vaccination.

## Main Text

Flaviviruses are a group of medically important viruses, including Zika (ZIKV), dengue (DENV), and West Nile (WNV) viruses that have recently circulated in the Western Hemisphere. Immature flavivirus virions bud into the endoplasmic reticulum (ER). They are composed of an ER-derived lipid bilayer, a nucleocapsid core containing multiple copies of the capsid (C) structural protein, an RNA genome, and 180 copies each of the structural proteins premembrane (prM) and envelope (E) incorporated into the viral membrane as heterotrimeric spikes (*1*). prM prevents the adventitious fusion of immature virions during egress through acidic compartments of the secretory pathway. During the transit of immature virions through the low pH environment of the trans-Golgi network, rearrangement of E and prM exposes a furin cleavage site in prM (*2*). Cleavage of prM has been shown to be required for the transition to an infectious mature form of the virus (*3*). The 94 amino acid pr cleavage product remains non-covalently associated with the virion until released from the cell (*2, 4*).

The E proteins of mature virions are organized as 90 sets of anti-parallel homodimers and are a principal target of neutralizing antibodies (*5*). The short membrane-bound M peptide is located beneath the E protein herringbone lattice. Neutralizing antibodies recognize E protein epitopes within a single protomer and surfaces extending across E dimers (*6, 7*). Flavivirus prM cleavage can be inefficient (*8, 9*). Antibody recognition differs among the immature and mature surfaces retained on partially mature particles (*7, 10, 11*). A capacity to neutralize mature virions correlates with vaccine-elicited protection from ZIKV infection in nonhuman primates (*12*).

The spread of ZIKV to the Western Hemisphere in 2015 prompted the rapid development of ZIKV vaccines. Numerous candidates were engineered to express the ZIKV structural proteins prM and E (*13*), including VRC5283, a DNA vaccine shown to be protective in animal models and immunogenic in Phase I clinical trials (*14, 15*). Co-expression of prM and E is sufficient to drive the production of non-infectious subviral particles (SVPs) lacking a nucleocapsid core (*16, 17*). SVP antigens have been explored as vaccines for multiple flaviviruses (*18, 19*) (*20*), with several candidates progressing to clinical trials (*21-23*). Curiously, an adenovirus vaccine expressing the ZIKV E protein preceded by a truncated prM (M-E) was protective in animal models and immunogenic in preclinical studies and clinical trials. However, the structure of this vaccine antigen was puzzling (*24, 25*). The position of the amino-terminal pr portion of prM within the heterotrimeric spikes of newly formed immature virions and the proposed roles of prM during the trafficking of virions suggested that particle formation may not occur without pr, or that virus-like particles produced following M-E expression may fuse with cellular membranes before release (*26, 27*). Consistent with this, prior Western blot studies of M-E transfected cells failed to detect extracellular antigens (*28*). Thus, the antigenic structure of an M-E immunogen has remained elusive.

### ZIKV subviral particles form in the absence of pr

To investigate the antigenic structure of M-E vaccine antigens, we established inducible CHO cell lines that uniformly expressed either ZIKV prM-E or M-E and performed transmission electron microscopy (**Fig. 1A**, and **fig. S1A**). In prM-E expressing cells, we observed round SVPs with an approximate diameter of ∼30 nm (**Fig. 1, B and C**), suggestive of an arrangement of 30 E dimers with T=1 icosahedral symmetry, as described for tick-borne encephalitis virus (TBEV) (*16, 29*). Notably, an abundance of SVPs was observed in M-E expressing cells, despite the absence of pr (**Fig. 1B**). M-E SVPs were larger (∼56 nm in diameter) than those produced following prM-E expression (**Fig. 1C**), and similar in diameter to infectious ZIKV virions that contain a nucleocapsid and display 90 E dimers on the surface (*30*). While individual M-E SVPs were found dispersed in membrane-bound structures throughout the cell, many were packed into large, membrane-bound structures resembling stacked sheets and lattices (**fig. S2, fig. S3, movie S1**). These data indicate that the pr portion of prM is dispensable for virion assembly and budding into the ER. Their accumulation suggests pr may play a role in virus particle trafficking. In support of a trafficking role for pr, ZIKV E protein release into supernatants of M-E expressing cells was delayed compared to cells expressing prM-E (**Fig. 1D**, and **fig. S1B**). A similar pattern was observed in a transient transfection model of ZIKV prM-E and M-E using 293T cells (**fig. S4**).

**Fig. 1.**
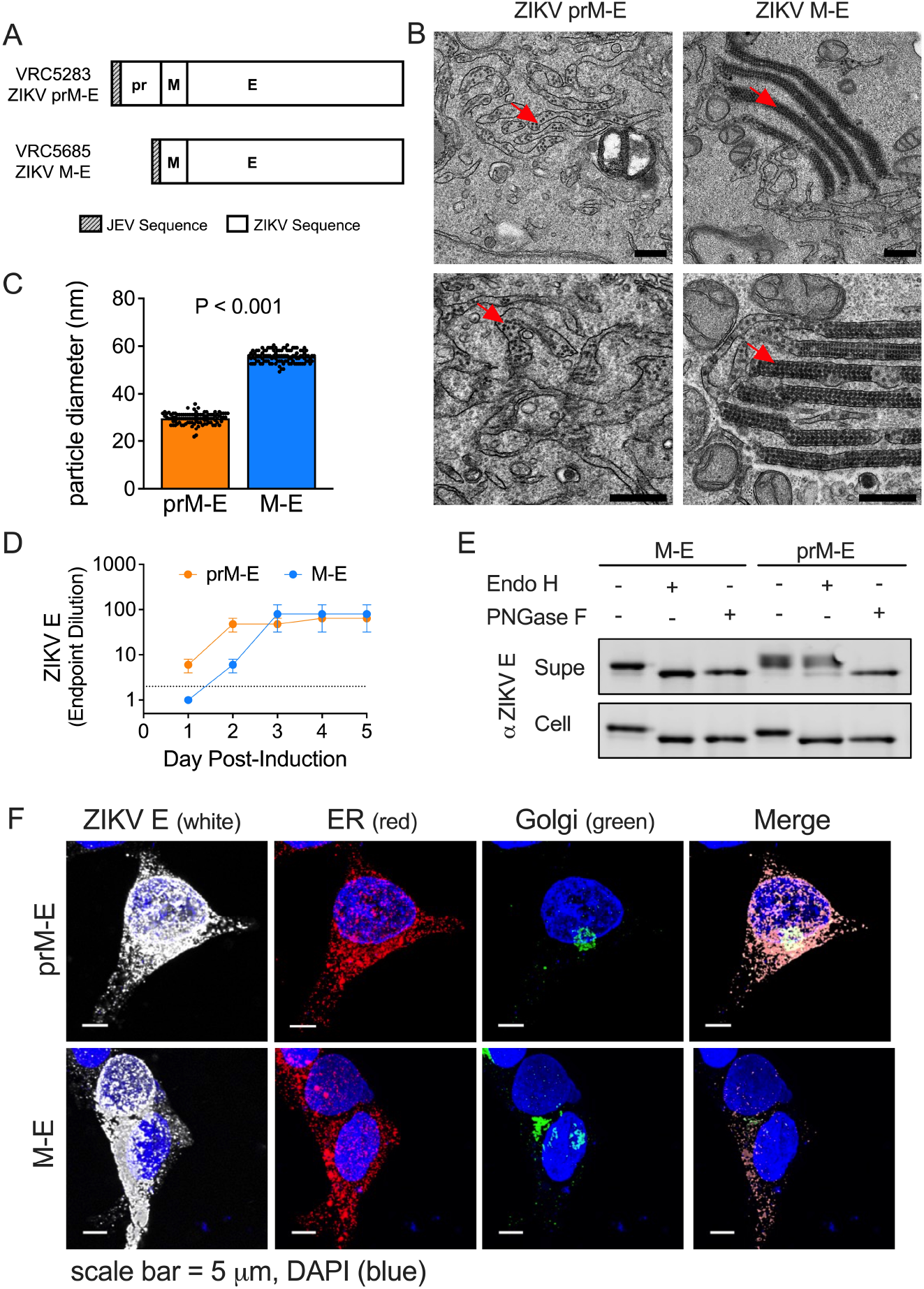
pr-independent biogenesis of ZIKV subviral particles. (**A**) DNA plasmids VRC5283 and VRC5685. (**B**) Transmission electron microscopy images of ZIKV SVPs in tetracycline-induced CHO cells 2 days post-induction. Red arrows indicate SVPs. Scale bars represent 0.5 microns. (**C**) Average SVP diameter calculated from 101 and 105 individual particles from prM-E and M-E images, respectively. Error bars indicate the standard error. (**D**) ZIKV E protein ELISA endpoint dilutions for supernatants sampled from tetracycline-induced CHO cell lines incubated at 37°C. Data is the average of two independent experiments. Error bars indicate the range. Samples that were not positive at an initial 1:2 dilution (dotted line) were assigned an endpoint dilution of 1. (**E**) M-E and prM-E ZIKV SVPs collected from the supernatant (Supe) or from inside transfected 293T cells (Cell) were left untreated or incubated with Endo H or PNGase F glycosidases, followed by SDS-PAGE and Western blotting with a ZIKV E protein-specific mAb. (**F**) Confocal microscopy of 293T cells transfected with ZIKV prM-E or M-E expressing plasmids. Transfected cells were fixed, and stained with ER, Golgi, and ZIKV E protein-specific antibodies. Scale bars represent micrometers.

To investigate differences in the intracellular trafficking and release of M-E particles, we monitored ZIKV E protein asparagine-linked glycan processing in transfected 293T cells. prM-E and M-E SVPs were collected from the supernatant or from cell lysates and treated with the glycosidase PNGase F (which cleaves all complex and hybrid N-linked glycans), or Endo H (which is specific for mannose-rich sugars and is unable to cleave sugars once processed by alpha-mannosidase II in the Golgi). Glycosidase treatment was followed by SDS-PAGE and Western blotting with a ZIKV E-specific monoclonal antibody (mAb) (**Fig. 1E**). PNGase F removed the E glycan from all SVPs, regardless of the construct (prM-E versus M-E) or location of harvest (supernatant versus cell lysate). In contrast, Endo H treatment yielded distinct patterns depending on the harvest site. The sugars of cell-associated prM-E and M-E SVPs were sensitive to Endo H cleavage, suggesting that most virus-like particles inside transfected cells had not trafficked through the Golgi. prM-E SVPs collected from the supernatant were Endo H resistant, indicating conventional trafficking through the secretory pathway. Interestingly, M-E SVPs collected from the supernatant remained sensitive to Endo H, suggesting virion exit without traveling through the Golgi compartment. In support, confocal microscopy studies revealed reduced E signal colocalization with a Golgi marker in cells transfected with ZIKV M-E as compared to ZIKV prM-E (**Fig. 1F**). Our results suggest that without pr, ZIKV M-E particles accumulate in the cell but do not efficiently traffic through the Golgi. We hypothesize that the M-E SVPs we detected in the supernatant are released after incidental cell lysis during culture.

### pr is not required for the production of an infectious flavivirus particle

To explore whether the E proteins arranged on M-E virus particles were functional, we produced reporter virus particles (RVPs) by complementation of ZIKV M-E with plasmids encoding ZIKV C and a GFP-expressing WNV subgenomic replicon (*31, 32*). Supernatants were sampled longitudinally from transfected 293T cells and assayed for infectious RVPs (**Fig. 2A**) and E protein content (**fig. S5**). Infectious M-E RVPs were detected starting at day 4 post-transfection, albeit at low titers when compared to standard RVPs produced in parallel with a prM-E plasmid. Despite the low titer, the specific infectivity (infectious titer/viral RNA) of ZIKV M-E RVPs was higher than standard RVPs (**Fig. 2B**). Mechanical cell lysis revealed that the majority of infectious M-E RVPs were retained intracellularly, whereas the prM-E RVPs were secreted into the supernatant (**Fig. 2, C and E**). As observed for SVPs, ZIKV M-E RVPs collected from the supernatant remained sensitive to Endo H (**Fig. 2F**). Complementation studies performed with M-E sequences from DENV serotype 2 (DENV2) and WNV also generated infectious M-E RVPs that accumulated within cells, establishing that pr-independent production of virus particles is possible for other medically important flavivirus species (**Fig. 2, G and H**).

**Fig. 2.**
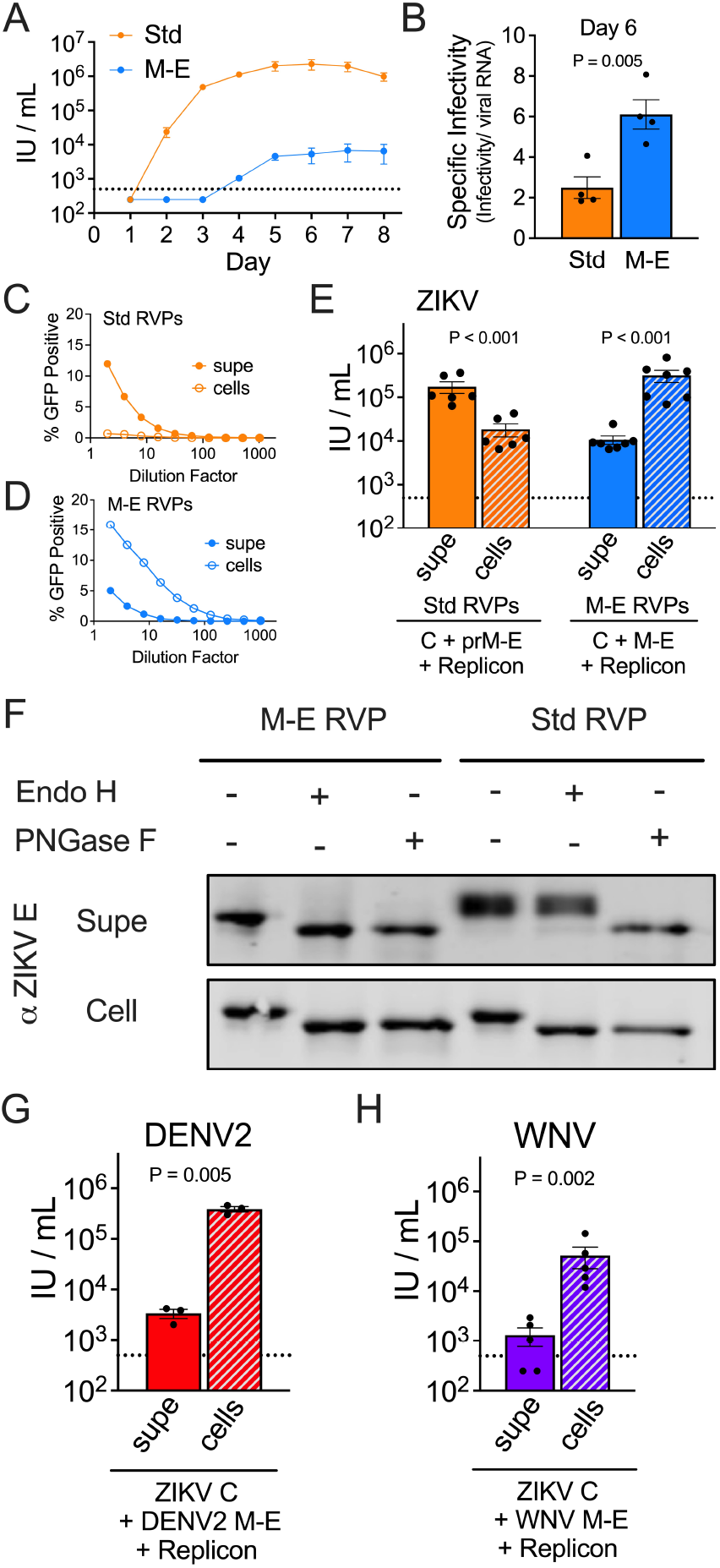
Infectious ZIKV M-E virus accumulates in cells. RVPs were generated in 293T cells with plasmids encoding ZIKV prM-E (Standard [Std]) or M-E. (**A**) RVP infectious titers in the supernatant were assessed longitudinally on Raji-DCSIGNR cells. Data are the average of two independent experiments. Error bars indicate the range. (**B**) RVP specific infectivity from 4 independent transfections. (**C-E**) ZIKV RVP infectivity in the supernatant at day 6 was compared to intracellular RVPs adjusted to an equivalent volume. (**C-D**) Representative RVP infectivity dose-response curves. Error bars indicate the range of duplicate infections. (**E**) Infectious titers calculated from 6-7 independent transfection experiments. Error bars indicate the standard error. **(F)** ZIKV RVPs from the supernatant or from lysed cells were incubated with Endo H or PNGase F glycosidases, followed by SDS-PAGE and Western blotting with a ZIKV E protein-specific antibody. (**G**) DENV2 M-E RVP infectious titers at day 6 from 3 independent experiments. Error bars indicate the standard error. (**H**) WNV M-E RVP infectious titers at day 6 from 5 independent experiments. Error bars indicate the standard error. The dotted lines indicate the LOD of the assay; titers below this threshold were assigned a value of half the LOD.

### A role for ZIKV pr in virion egress

The N-linked glycan at pr residue 70 is required to produce infectious ZIKV virions (*33*). We hypothesized that mutations abolishing the N-linked glycan on pr would yield RVPs with a similar trafficking phenotype as M-E RVPs. RVPs were generated using prM-E plasmids with an alanine mutation incorporated at prM residue 71 or 72, both threonines in the wild type sequence (**Fig. 3**). Mutation at residue 72, but not 71, abolishes the N-linked glycan motif N-X-S/T, where the T at the +2 position, but not position +1 is critical for glycan addition. The presence or absence of the prM N-linked glycan was confirmed by Western blotting of PNGase F-treated prM T71A and T72A RVP variants with a pr-specific mAb (**Fig. 3B**). We next repeated transfection experiments to compare the infectivity of secreted versus intracellular virus (**Fig. 3C**). Infectious prM T72A RVPs, but not wild type or the T71A variant RVPs, were more abundant inside cells than in the supernatant. Limited quantities of the prM T72A variant RVPs were also observed by measuring the E protein content in the supernatant (**Fig. 3D**). Of note, measures of E protein content include both infectious mature virions and non-infectious immature virions. Thus, the high levels of E protein detected inside cells producing wild type and prM T71A RVPs reflects the low specific infectivity of these virus populations. These data identify the pr N-linked glycan as a required component of virion trafficking.

**Fig. 3.**
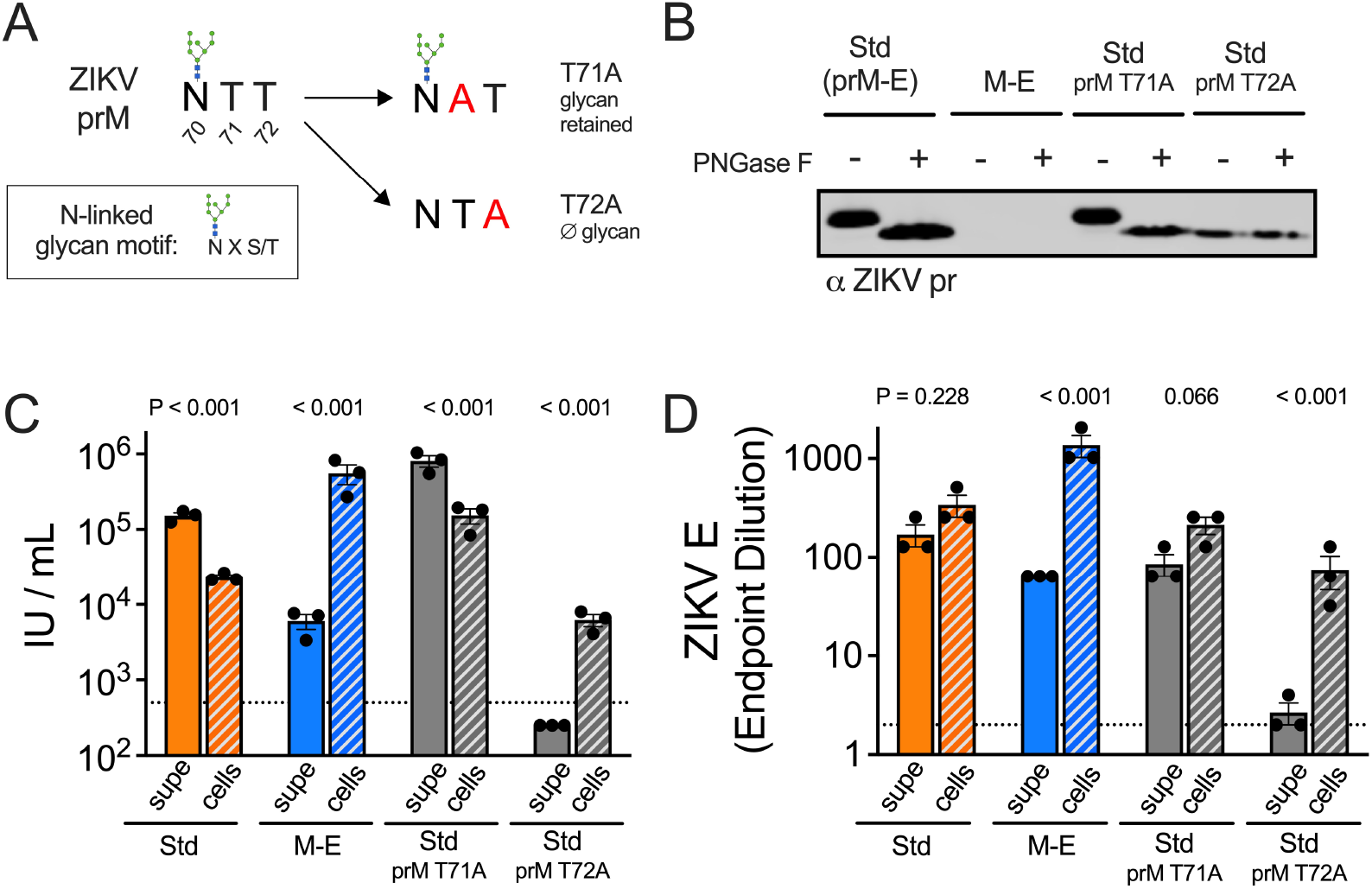
ZIKV pr is important for virion egress from cells. (**A**) Diagram of the N-linked glycan sequence motif in wild type or mutated prM. (**B**) ZIKV prM-E plasmids with mutations at prM T71A or T72A were used to generate RVPs. Wild type or variant RVPs collected from the supernatant were left untreated or incubated with PNGase F, followed by SDS-PAGE and western blotting with a ZIKV pr-specific antibody. The infectivity (**C**) and E protein ELISA endpoint dilution (**D**) of the indicated ZIKV RVPs released into the supernatant at day 6 post-transfection was compared to intracellular RVPs obtained via multiple freeze-thaw cycles and adjusted to an equivalent volume. Shown are data from 3 independent experiments. Bar graphs and error bars indicate the average and standard error. Dotted lines indicate the LOD of the assays, a titer of 500 IU/mL or the ELISA starting dilution of 2. Results below these thresholds were assigned a value of half the LOD.

### M-E virus-like particles retain antigenic features of mature virions

The extent of maturation impacts the antigenic structure of infectious virions and SVPs and may shape the antibody repertoire elicited by infection and vaccination. Neutralizing antibodies targeting the mature form of the ZIKV virion correlated with protection from infection following VRC5283 prM-E vaccination (*12*). To probe the antigenic structure and maturation state of virions formed in the absence of an intact prM protein, neutralization studies of ZIKV M-E RVPs were performed using well-characterized mAbs (**Fig. 4, A, B, C, D, E and F**). As anticipated, M-E RVPs were completely resistant to neutralization by the pr-reactive mAb 19pr, as were wild type RVPs produced in cells over-expressing furin (+Furin) (**Fig. 4A**). In contrast, neutralization of standard RVP preparations revealed the expected sigmoidal-shaped dose-response curve and resistant fraction due to the presence of partially mature particles that retain uncleaved prM (*34*). ZIKV M-E RVPs were efficiently neutralized by an E-dimer epitope (EDE)-specific mAb, demonstrating an antigenic surface similar to infectious ZIKV (**Fig. 4B**). Similarly, DENV2 and WNV M-E RVPs were resistant to mAbs that neutralize infection in a maturation-state sensitive manner (**Fig. 4, C and E**), yet highly sensitive to maturation-state insensitive neutralizing mAbs (**Fig. 4, D and F**).

**Fig. 4.**
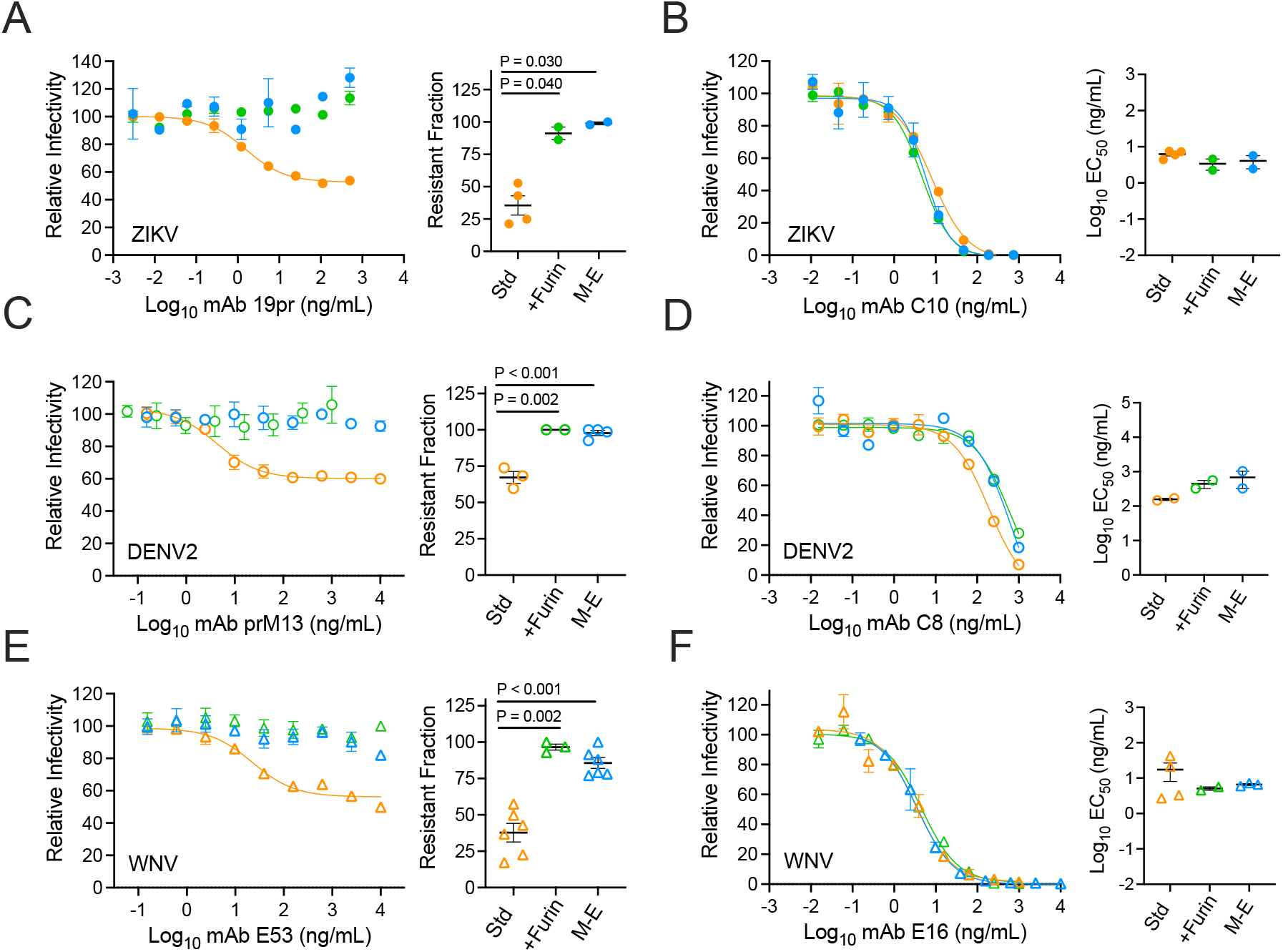
Antigenicity of flavivirus M-E virus. Representative dose-response neutralization profiles for ZIKV, DENV2 or WNV RVPs (Std, +Furin, and M-E) with the indicated mAbs are shown on the left. Error bars indicate the range of duplicate infections. EC_50_ or resistant fraction (the % not neutralized at highest mAb concentration) from 2-6 independent experiments are shown on the right. Horizontal line and error bars indicate the mean and standard error. (**A, C, E**) Neutralization with mAbs sensitive to virus maturation. mAbs 19pr and prM13 bind to pr of ZIKV and DENV2, respectively (*34*). The WNV-specific mAb E53 binds an E DII fusion loop epitope that is accessible on an immature virion but buried on the surface of the mature virion (*10*). (**B, D, F**) Neutralization with mAbs that are not sensitive to the extent of virus maturation. mAbs C8 and C10 bind E-dimer epitopes and are cross-reactive against ZIKV and DENV (*6*). The WNV-specific mAb E16 binds an epitope on the lateral ridge of E DIII (*52*). Groups in A-F were compared using a one-way ANOVA; only P values < 0.05 are shown.

### Immunogenicity of a ZIKV M-E DNA vaccine

DNA and adenovirus vaccines encoding ZIKV M-E antigens are protective in animal models (*25, 28*). Our data suggested that the M-E SVPs produced by these vaccines retain key antigenic features of mature virions and are of similar size. To explore how the unique biogenesis of this antigen shapes the antibody response relative to prM-E vaccines, VRC5685 DNA expressing ZIKV M-E was administered intramuscularly to rhesus macaques in two doses, 4 weeks apart using a needle-free device. Groups of four animals received 1, 0.3, or 0.1 mg doses at weeks 0 and 4. Neutralizing antibody titers against standard ZIKV RVPs were assessed at weeks 0, 6, and 8 after initial immunization (**Fig. 5A**). VRC5685 vaccination elicited neutralizing antibodies in all animals (reciprocal EC_50_ titer ranging from 64-1969 at week 8). All animals were protected from viremia following challenge with ZIKV strain PRVABC59 (**Fig. 5B**). Similar neutralizing antibody titers and protection were observed in animals vaccinated with VRC5283 (encoding prM-E) and challenged (*12*).

**Fig. 5.**
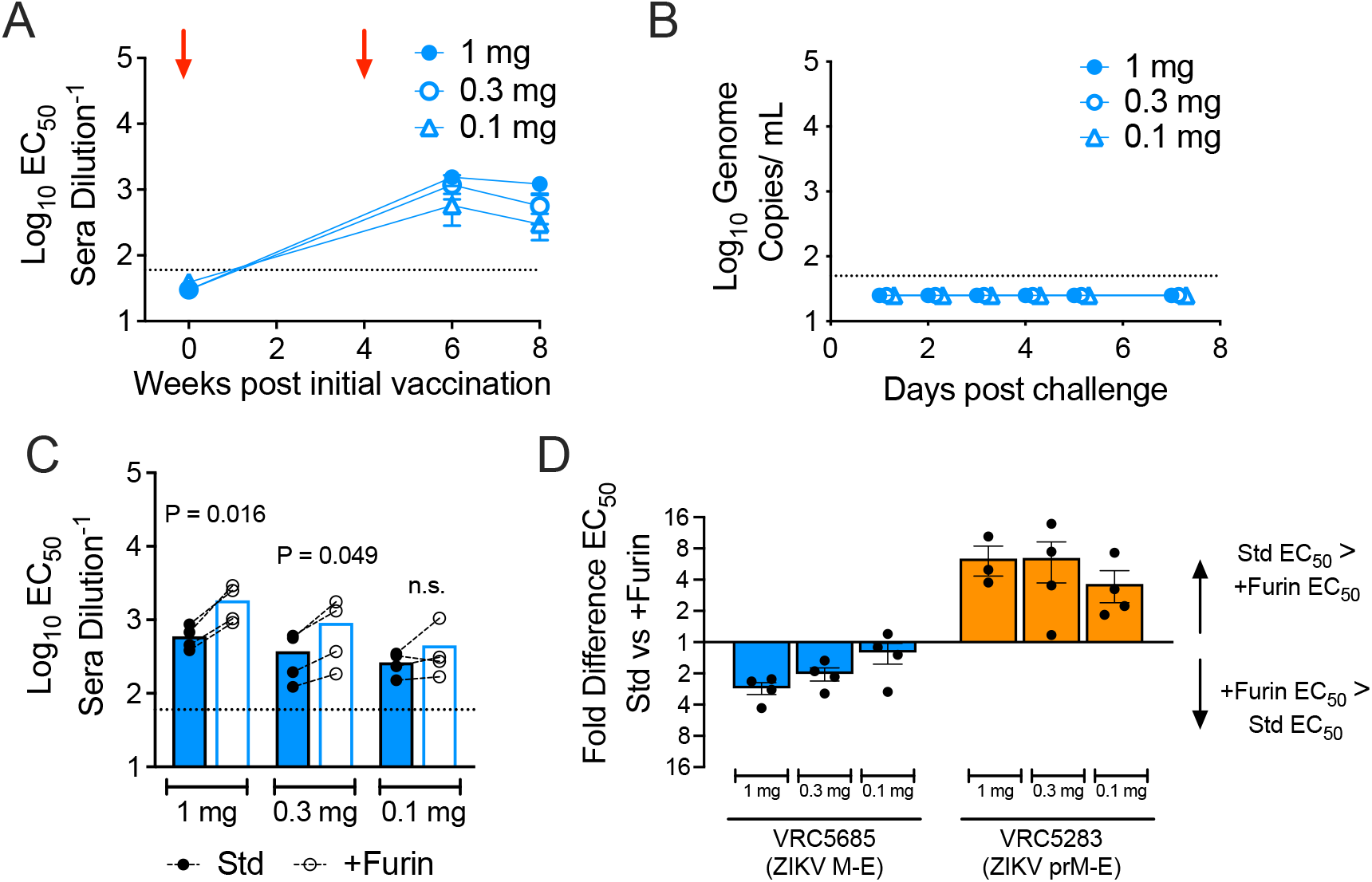
Immunogenicity of a ZIKV M-E vaccine. Macaques (four per group) were immunized at weeks 0 and 4 with 1, 0.3, or 0.1 mg of DNA plasmid VRC5685, and challenged with ZIKV at week 8. (**A**) Serum were used in neutralization assays with standard ZIKV RVPs. Dots represents the average EC_50_ titer for each dose group; error bars indicate the standard error. Red arrows indicate administration of vaccine. (**B**) Viral loads were measured by qRT-PCR after challenge with ZIKV. Each dot represents the average viral load for each dose group. (**C**) Week 8 serum was used in pairwise neutralization assays against standard and +Furin ZIKV RVPs. Each dot represents the average of three experiments for an individual animal. Dashed lines connect EC_50_ titers from the same animal. Bars denote the group average. Std vs +Furin EC_50_ values per dose group were compared by a one-way ANOVA. Dotted lines represent the LOD of the assay. (**D**) The data from panel C is expressed as the fold difference of Std vs +Furin EC_50_ values per dose group and shown in comparison to previously published data from animals that had received the ZIKV prM-E DNA vaccine candidate VRC5283 (*12*).

To characterize the quality of the antibody response elicited by VRC5685 vaccination, we performed pairwise comparisons of the ability of macaque sera to neutralize standard and more homogeneously mature (+Furin) ZIKV RVP preparations. Neutralization assays using sera collected at week 8 (Fig. 5C) indicated that VRC5685 vaccination elicited a neutralizing antibody response that was largely insensitive to the maturation state of the virus. Most individual macaque sera neutralized +Furin RVPs slightly better than standard RVPs, and this difference was significant for the 1 mg and 0.3 mg dose groups. In contrast, our prior studies revealed that vaccination with plasmids expressing prM-E elicited neutralizing antibodies that were less efficient at neutralizing mature forms of the virion lacking uncleaved prM (**Fig. 5D**; published VRC5283 titers are included to simplify comparisons)(*12*). Overall, these results indicate that the ZIKV M-E DNA plasmid VRC5685 provides protection from ZIKV infection and elicits a neutralizing antibody response that is insensitive to the maturation state of the virus, making it qualitatively better than a similar prM-E vaccine.

The mechanisms of flavivirus assembly and budding are not well understood. The expression of prM and E is sufficient to produce SVPs capable of catalyzing membrane fusion. Immature particles display a “spiky” surface comprised of prM_3_:E_3_ heterotrimers (*35*). Trimer formation has been hypothesized to position the transmembrane domains of prM and E to induce the membrane curvature required for budding into the ER (*26*), however, experimental evidence supporting this model is limited. Here, we demonstrate that flaviviruses can assemble without pr. It is unclear whether the E proteins incorporated into M-E virus particles form dimers immediately after synthesis or if a trimeric state is possible without pr at the distal end. The larger diameter of ZIKV M-E SVPs suggests the lack of pr influences membrane curvature during budding, perhaps due to differences in the arrangement of transmembrane domains in the absence of stable trimers. Recently, the spike architecture of immature flaviviruses was resolved at a higher resolution, allowing refinement of the molecular interactions between prM and E (*36, 37*). The proposed “ripcord” model for the rearrangement of E trimers to dimers arising from these structures postulates that when prM:E interactions are disrupted by exposure to low pH in the Golgi, the E proteins simply collapse (*36, 38*). Thus, in the absence of interactions between pr and E, trimer formation may be impossible, allowing M-E antigens to directly form dimers.

Our data show that M-E particles readily bud into the ER, but these particles are released from cells inefficiently and do not traffic through the Golgi. These observations suggest that pr functions to chaperone virions during egress. Further, we demonstrate that removing the N-linked glycan of pr is sufficient to limit the transit of virus particles from the ER to the Golgi, in agreement with prior studies of WNV and Japanese encephalitis virus (JEV) (*39, 40*). Indeed, flavivirus pr has been hypothesized to originate from a cellular HSP40-like chaperone gene that eventually evolved into an accessory protein required to navigate low-pH exposure during viral egress (*41*). The infectivity of M-E RVPs establishes that the E proteins of M-E virions are arranged to support viral attachment and membrane fusion. prM prevents oligomeric changes in the E protein at low pH during egress (*4*). Without this protection, M-E virions not retained in the ER would be expected to undergo adventitious fusion during transit through acidic compartments of the Golgi.

Recent studies highlight the importance of antibody quality in flavivirus immunity (*12, 42*). Immunization studies with two similar ZIKV prM-E DNA vaccine candidates identified an ability to neutralize mature forms of the virus as a surrogate of protection (*12*). Complex quaternary surfaces across the E dimers of mature virions contain epitopes recognized by neutralizing mAbs, including E dimer epitopes (EDE) originally described for DENV. E dimers are recognized by polyclonal immune sera against DENV, yellow fever virus, ZIKV, and TBEV (*43-46*). Conversely, antibodies that bind epitopes available on the prM_3_:E_3_ trimers, including those that bind pr, may be immunodominant, cross-reactive, and possess limited neutralization potential (*34, 47-49*). Reducing the elicitation of poor-quality antibodies is an important goal of flavivirus vaccine development, as they are linked to poor prognosis following some secondary infections (*5, 50, 51*).

SVP antigens have shown promise for several flaviviruses when expressed via multiple vaccine platforms, including mRNA. An incomplete understanding of the structure of these heterogeneous virus particles limits structure-guided rational design efforts. Pre-clinical and clinical studies demonstrated the safety and immunogenicity of an adenovirus-vectored ZIKV M-E candidate (*24, 25*). Here, we establish that the unique characteristics of M-E antigens favor a high-quality neutralizing antibody response. M-E particles do not incorporate the prM_3_:E_3_ trimer spikes of partially mature or immature virions, nor can they elicit poorly neutralizing pr-specific antibodies (*34, 48*). While we do not yet know the structure of virus particles assembled without pr, M-E SVPs were of similar size as an infectious virion. Differences in the arrangement of E proteins on the smaller SVPs resulting from prM-E expression elicits a distinct antibody response (*16, 17, 29*). The unexpected larger size of M-E SVPs may better recapitulate the E protein rafts that comprise the surface of infectious virions. Serum from M-E immunized nonhuman primates efficiently neutralized mature forms of ZIKV, in contrast to serum elicited against prM-E vaccine candidates of similar sequence. Critically, these data establish that M-E antigens elicited a higher-quality antibody response. That we were able to produce M-E virions with other flaviviruses identifies this as a potential general solution to flavivirus vaccination.

## Materials and Methods

### Ethics statement

This study was carried out in accordance with the recommendations and guidelines of the National Institutes of Health (NIH) Guide to the Care and Use of Laboratory Animals. The protocol was approved by the Animal Care and Use Committee of the Vaccine Research Center (VRC), National Institute of Allergy and Infectious Diseases (NIAID) at the NIH. Rhesus macaques (Macaca mulatta) were housed and cared for in accordance with local, state, federal, and institutional policies at BIOQUAL Inc., an Association for Assessment and Accreditation of Laboratory Animal Care International (AAALAC) accredited facility.

### Cell lines

HEK-293T/17 cells were maintained in in Dulbecco’s modified Eagle medium (DMEM) with GlutaMAX (ThermoFisher Scientific) supplemented with 7% fetal bovine serum (FBS) and 100 U/ml penicillin streptomycin (PS). Raji B lymphocytes expressing DC-SIGNR (Raji-DCSIGNR) were cultured in RPMI-1640 medium (ThermoFisher Scientific) supplemented with 7% FBS and 100 U/ml PS (*53*). Parental T-REx–Chinese hamster ovary (CHO) cells were maintained in F12 media supplemented with 7% FBS and 0.01 mg/ml Blasticidin S HCl. All cells were maintained at 37°C and 7% CO_2_.

### Plasmids

Plasmid VRC5685 (ZIKV M-E) was generated by deleting the pr portion of prM from the previously described plasmid VRC5283 that encodes ZIKV prM-E (strain H/PF/2013) (*14*). At the 5’ end of the open reading frame, both plasmids share the amino acid sequence MGKRSAGSIMWLASLAVVIACAGA, which encodes the prM signal sequence from JEV (*54*). Immediately following this sequence, the next six amino acids of VRC5283 versus VRC5685 are AEVTRR and AVTLPS, respectively. Both sequences were cloned into the previously described mammalian expression plasmid VRC8400 (*14, 23, 55*). Sequences encoding prM-E and M-E from DENV2 strain 16681 and WNV strain NY99 were similarly cloned downstream of the JEV signal sequence into the VRC8400 vector.

Additional plasmids used for RVP production (subgenomic replicon pWNVII-Rep-GFP/Zeo, C-prM-E structural gene plasmids for ZIKV, DENV2 and WNV, human furin) have been previously described (*31, 32, 56*). A plasmid encoding ZIKV C was synthesized by Genscript and cloned into pcDNA3.1(+).

### Non-human primate studies

DNA plasmid VRC5685, encoding ZIKV M-E from strain H/PF/2013, was included in a previously described dose de-escalation study of ZIKV prM-E vaccine candidates VRC5283 and VRC5288 in rhesus macaques (*12*). Groups of four animals received de-escalating doses of VRC5283, VRC5288, or VRC5685 DNA (1, 0.3, and 0.1 mg), or 1 mg of the empty vector VRC8400 as a control. All animals received split-dose injections of the respective vaccine in each quadricep (for example, two 0.5 mg injections for the 1 mg dose group) at weeks 0 and 4 using a needle-free delivery system (PharmaJet). Animals were challenged at week 8 with 10^3^ FFU of ZIKV PRVABC59 administered subcutaneously. Animals were bled at various timepoints for quantification of viral loads and serum neutralizing antibody titers. Data obtained from VRC5283, VRC5288, and VRC8400 vaccinated animals were published in a prior study (*12*). The current study reflects the first report of the VRC5685 vaccinated group, with neutralization data from the previously published VRC5283 group included for comparison.

### ZIKV SVP production in 293T cells

HEK-293T/17 cells pre-plated in 6-well dishes were transfected with 3μg VRC5283 (prM-E), VRC5685 (M-E) or empty vector control (VRC8400) plasmids using Lipofectamine 3000 according to the manufacturer’s instructions. Cells were incubated at either 30°C or 37°C and supernatants were passed through a 0.2 μm filter and stored at -80°C until use.

### ZIKV prM-E and M-E tetracycline-inducible cell lines

A tetracycline-inducible cell line expressing ZIKV prM-E has been previously described (*12*). An additional cell line expressing ZIKV M-E was generated using the same methods. Briefly, the open reading frame from plasmid VRC5685, encoding the JEV signal sequence upstream of ZIKV M-E, was PCR-amplified and cloned into the pT-RExDEST30 mammalian expression vector using Gateway technology (Invitrogen) to generate the plasmid pT-REx-ZIKV M-E. Parental T-REx–Chinese hamster ovary (CHO) cells maintained in F12 medium supplemented with 7% FBS and blasticidin S HCl (0.01 mg/ml; Gibco) were transfected with the pT-REx-ZIKV M-E plasmid using Lipofectamine 3000 (Invitrogen). The following day, medium was replaced with F12 medium supplemented with 7% FBS, blasticidin S HCl (0.01 mg/ml), and Geneticin (0.5 mg/ml; Gibco) to select for transfected cells. After confirming expression in the resulting polyclonal cell line, cells were sufficiently diluted in 96-well plates to achieve a single cell per well, and monoclonal cell lines were expanded in F12 medium supplemented with 7% FBS, blasticidin S HCl (0.01 mg/ml), and Geneticin (0.5 mg/ml). To induce ZIKV structural protein expression, cells were induced with tetracycline (500 ng/ml) and maintained in F12 medium supplemented with 1% FBS, blasticidin S HCl (0.01 mg/ml), and Geneticin (0.5 mg/ml) at 37°C. Parental T-REx–CHO cells were induced with tetracycline (500 ng/ml) and maintained in F12 medium supplemented with 1% FBS and blasticidin S HCl (0.01 mg/ml) at 37°C as a negative control. Supernatants were filtered through a 0.2 μm membrane and stored at -80°C until use.

### ELISA

ZIKV E protein was detected using a previously described particle-capture ELISA format (*12*). 384-well microplates (Corning) were coated with 3 µg/ml of the E-specific mAb ZV-67 in 0.1 M carbonate-bicarbonate buffer, pH 9.4 (ThermoFisher Scientific) and incubated at 4°C overnight. Plates were blocked with 5% skim milk in phosphate-buffered saline, 0.1% Tween (PBST) at 37°C for 1 hour, followed by a single wash with PBST. Supernatants serially diluted in PBST + 5% skim milk were added to plates, incubated at 37°C for 1 hour, followed by six washes with PBST. Biotinylated ZV-67 (DSB-X Biotin protein labeling kit, ThermoFisher) was added to plates and incubated at 37°C for 1 hour, followed by six washes with PBST. High sensitivity streptavidin-HRP (ThermoFisher Scientific) was added and incubated at 37°C for 1 hour, followed by six washes with PBST. The assay was developed using 3,3’,5,5’-tetramethylbenzidine (TMB) (SeraCare) followed by 1N HCl and the absorbance measured at 450 nm (BioTek Synergy H1). The endpoint dilution was calculated as the final dilution > 2-fold above the average background OD450. To detect uncleaved prM protein, a modified version of this ELISA was performed by replacing mAb ZV-67 with the ZIKV pr-specific mAb 19pr (generated in-house).

### Viral RNA Quantification by Reverse Transcription Polymerase Chain Reaction

ZIKV RNA was quantified for the rhesus macaque study by BIOQUAL Inc. using qRT-PCR as previously described (*12*). Standard curves were generated using serial dilutions of viral RNA with a known concentration. The limit of detection (LOD) was determined to be 50 genome copies/ml. Values are the average of one to three technical replicates.

Genomic viral RNA in RVP samples was quantified using a previously described assay (*57, 58*). Standard curves were generated using known copy numbers of the WNV subgenomic replicon plasmid. Samples were run in triplicate, and potential plasmid DNA contamination in supernatants was accounted for by subtracting values obtained when the same sample was run in parallel without reverse transcriptase.

### Transmission Electron Microscopy Imaging

Control, ZIKV prM-E or ZIKV M-E CHO T-REx cells were seeded on 13mm round Thermanox coverslips (Electron Microscopy Sciences) in 24-well tissue culture plates. The next day, cells were induced by adding 1000 ng/mL tetracycline to the medium. On day 2 or 4 post-induction, cells on coverslips were fixed with 2.5% glutaraldehyde/ 2% paraformaldehyde in 0.1 M Sorenson’s sodium-potassium phosphate buffer, pH 7.2 (Electron Microscopy Sciences).

Samples were post-fixed with 1.0% osmium tetroxide/ 0.8% potassium ferricyanide in 0.1 M sodium cacodylate buffer, washed with buffer then stained with 1% tannic acid in dH_2_O. After additional buffer washes, the samples were further osmicated with 2% osmium tetroxide in 0.1 M sodium cacodylate, then washed with dH_2_O and additionally stained overnight with 1% uranyl acetate at 4° C. After 3 × 5 minute dH_2_O washes, the samples were dehydrated with a graded ethanol series, and embedded in Spurr’s resin. Thin sections were cut with a Leica UC7 ultramicrotome (Buffalo Grove, IL) prior to viewing at 120 kV on a Thermo Fisher/ FEI BT Tecnai transmission electron microscope (Thermo Fisher/ FEI, Hillsboro, OR) or Hitachi 7800 transmission electron microscope at 80kV (Tokyo, Japan). Digital images were acquired with a Gatan Rio camera (Gatan, Pleasanton, CA) on the Tecnai or the AMT XR81-B digital camera system on the Hitachi. Figures were assembled using Adobe Photoshop version 25.7.0 (Adobe, San Jose, CA).

### Electron Tomography

150 nm sections of samples prepared as above were placed on Formvar coated slot grids (Electron Microscopy Sciences, Hatfield, PA) and 10 nm colloidal gold (BBI Solutions, Portland, ME) was added as fiducial markers. Tilt-series were acquired using SerialEM (*59*) on a Tecnai T12 (Thermo Fisher, Waltham, MA) transmission electron microscope operating at 120 kV using a Rio CMOS digital camera (Model 1809, Gatan, Pleasanton, CA). Dual axis tilt-series (*60*) were collected +/-60 °, every 1° and tomograms were generated with IMOD (*61*) using weighted back projection. Segmentation and rendering were performed in Amira (Thermo Fisher, Waltham, MA). Movies were generated by rendering PNG images from Amira and assembling them in Adobe Premiere Pro (Adobe, San Jose, CA).

### Reporter Virus Particle Production

ZIKV RVPs were produced by complementation of a GFP-expressing WNV lineage II subgenomic replicon (pWNVII-Rep-GFP/Zeo) with plasmids encoding ZIKV strain H/PF/2013 structural proteins. Standard ZIKV RVPs were produced as previously described using a C-prM-E structural plasmid (*31*), and +Furin RVPs were produced by the inclusion of a plasmid encoding human furin (*12*). To generate RVPs with VRC5283 (prM-E) and VRC5685 (M-E) plasmids, a slightly modified strategy was used in which the ZIKV C protein was encoded from a separate plasmid, as has been described for WNV RVPs (*32, 62*). Using this three-plasmid strategy, HEK-293T/17 cells were co-transfected with the WNV subgenomic replicon, a plasmid encoding ZIKV C, and a plasmid encoding ZIKV prM-E or M-E at a ratio of 30:10:1 by mass, using Lipofectamine 3000 reagent. The decreased amount of prM-E or M-E plasmid limits the potential for SVP secretion. C-prM-E and M-E constructs were similarly used to make RVPs encoding the structural genes of DENV2 and WNV. Transfected cells were incubated at 30°C, and RVP-containing supernatants were passed through a 0.2 μm filter and stored at -80°C until use. When necessary for downstream applications, low titer RVP stocks were concentrated using an Amicon Ultra centrifugal filter with a 100 kDa molecular weight cut-off. To determine viral titers, two-fold serial dilutions of RVPs were used to infect Raji-DCSIGNR cells. Following incubation at 37°C for 2 days, cells were fixed with 2% PFA, and infected cells quantified by flow cytometry.

### Isolation of cell-associated RVPs

To isolate cell-associated RVPs, two distinct methods were used. <u>1) Freeze-thaw</u>: Transfected cells were washed with PBS, scraped to detach from the tissue culture plate, and transferred to a microcentrifuge tube. Three freeze-thaw cycles were performed in a -80°C freezer, followed by a 37°C thaw step. <u>2) Sonication</u>: Transfected cells were washed with PBS, scraped to detach from the tissue culture plate, and transferred to a microcentrifuge tube. Resuspended cells were sonicated three times at 70% amplitude for 1 min, separated by brief incubations on ice. Lysed cells were transferred to a microcentrifuge tube. For both methods, the cell suspension was subject to centrifugation at 2200xg for 10 min to remove cellular debris, passed through a 0.2 μm filter, and frozen at -80°C until use. Cell-associated RVPs were resuspended in a volume of media equivalent to the supernatant initially harvested.

### RVP neutralizing antibody assay

ZIKV RVP neutralization assays were performed as previously described (*31, 57*). Briefly, serial dilutions of mAbs or heat-inactivated macaque sera were incubated with RVPs for 1 hour at 37°C to allow for steady-state binding. Immune complexes were used to infect Raji-DCSIGNR cells and performed in technical duplicates. Infection was carried out at 37°C for 2 days, at which point cells were fixed at a final concentration of 2% paraformaldehyde, and GFP positive cells quantified by flow cytometry (FACSCelesta, Becton Dickinson). Non-linear regression analysis was performed to estimate the mAb concentration or reciprocal serum dilution required for half-maximal infectivity (EC_50_). In experiments performed with macaque sera, the limit of detection (LOD) was set as the reciprocal initial dilution of sera (1:60). EC_50_ titers estimated at a value <LOD were assigned a titer of one-half the LOD. When possible, RVP stocks were sufficiently diluted to ensure antibody excess at informative points of the dose-response curves. For ZIKV M-E RVP stocks that could not be diluted due to low infectivity, control mAbs with known EC_50_ values were included in neutralization assays to ensure that the percentage law was not in violation (*63*).

### ZIKV Structural Protein Immunoblotting

ZIKV E and prM proteins were detected by Western blotting. RVPs or SVPs were concentrated and purified by ultracentrifugation through a 30% sucrose cushion overnight at 28,000 rpm, 4°C in a Beckman SW32Ti rotor. Pelleted virus was resuspended in 1x PBS and E protein concentration estimated by ELISA. Concentrated virus was heated at 70°C for 10 minutes in LDS buffer (NuPAGE) and left untreated or treated with Endo H or PNGase F glycosidases following the manufacturer’s protocol (New England Biolabs). Samples (normalized by E protein signal) were loaded into a 1.5 mm, 4-12% Bis-Tris gel (NuPAGE) and run using SDS running buffer (NuPAGE) for 1 h at 150 V. Separated proteins were transferred to a nitrocellulose membrane (ThermoFisher) using an iBlot (ThermoFisher) at 25 V for 8 min. The membrane was blocked overnight at 4° C with Pierce blocking buffer (ThermoFisher), followed by incubation with biotinylated anti-ZIKV E (ZV-67) or biotinylated anti-ZIKV pr (19pr) mAb at 2 µg/mL for 1 h at RT. Membranes were then washed three times for 10 minutes each with PBST, followed by incubation with IRDye 800CW Streptavidin (LI-Cor) at a 1:2000 dilution for 1 h at RT. The membrane was washed again with PBST and the fluorescence signal imaged at 800 nm on an Odyssey CLx infrared imaging system (LI-Cor).

### Confocal Microscopy

293T cells were plated in Lab-Tek II chambered #1.5 coverglass 4 chamber slides (Nunc # 155382) (9×10^4^ cells per chamber in 0.5 mL medium). The next day, cells were transfected with 0.25 μg of ZIKV prM-E or M-E plasmid using Lipofectamine 3000 according to the manufacturer’s instructions. 16 hours later, cells were washed with PBS and then fixed with PBS + 2% paraformaldehyde for 15 minutes. Cells were then washed 2x with PBS, permeabilized with PBS + 0.1% TritonX-100 for 15 minutes, washed 2x with PBS, and blocked for 1 hour in 10% normal goat serum (ThermoFisher #50062Z). Cells were incubated for 1 hour with primary antibodies (1:1000 mouse Golgin-97 mAb clone CDF4 [Invitrogen #A-21270], 1:200 rabbit calreticulin polyclonal Ab [Invitrogen #PA5-25922], 1:200 ZV-67 mAb) diluted in blocking buffer. After 2x washes with PBS, cells were incubated for 1 hour with secondary antibodies (1:500 goat anti-human Alexa-fluor 647 [Invitrogen #A48279TR], 1:500 goat anti-rabbit Alexa-fluor 594 [Invitrogen #A32740], 1:500 donkey anti-mouse Alexa-fluor 488 [Invitrogen #A32766TR]). Cells were stained with DAPI, followed by 2x washes and storage in PBS. Chamber slides were imaged with a 63X objective on a Leica Stellaris microscope using Lightning deconvolution.

## Acknowledgments

We thank members of the Arbovirus Immunity Section, VRC, NIH for scientific advice and comments.

## Funding

This work from the Vaccine Research Center was funded by the Intramural Program of NIAID, NIH.

## Author contributions

Conceptualization: KAD, TCP

Methodology: KAD, MS, KEB, WS, LW, CLS, BTH, WPK, KMM, HDH, ERF

Investigation: KAD, MS, ES, BB, BMF, KEB, WS, LW, NC, DNG, CLS, BTH, MA, HDH

Funding acquisition: TCP

Supervision: KAD, BSG, ERF, TCP

Writing – original draft: KAD, TCP

Writing – review & editing: all authors

## Competing interests

The authors declare that they have no competing interests.

## Data and materials availability

All data are available in the main text or the supplementary materials. Reagents and remaining biospecimens are available from the NIH under material transfer agreements.

## List of Supplementary Materials

Figs. S1 to S5

Movie S1

## Notes

### Competing Interest Statement

The authors have declared no competing interest.

