## Supplementary Materials for "pr-independent biogenesis of infectious mature Zika virus particles"

**The PDF file includes:**

Figs. S1 to S5

**Other Supplementary Materials for this manuscript include the following:**

Movie S1

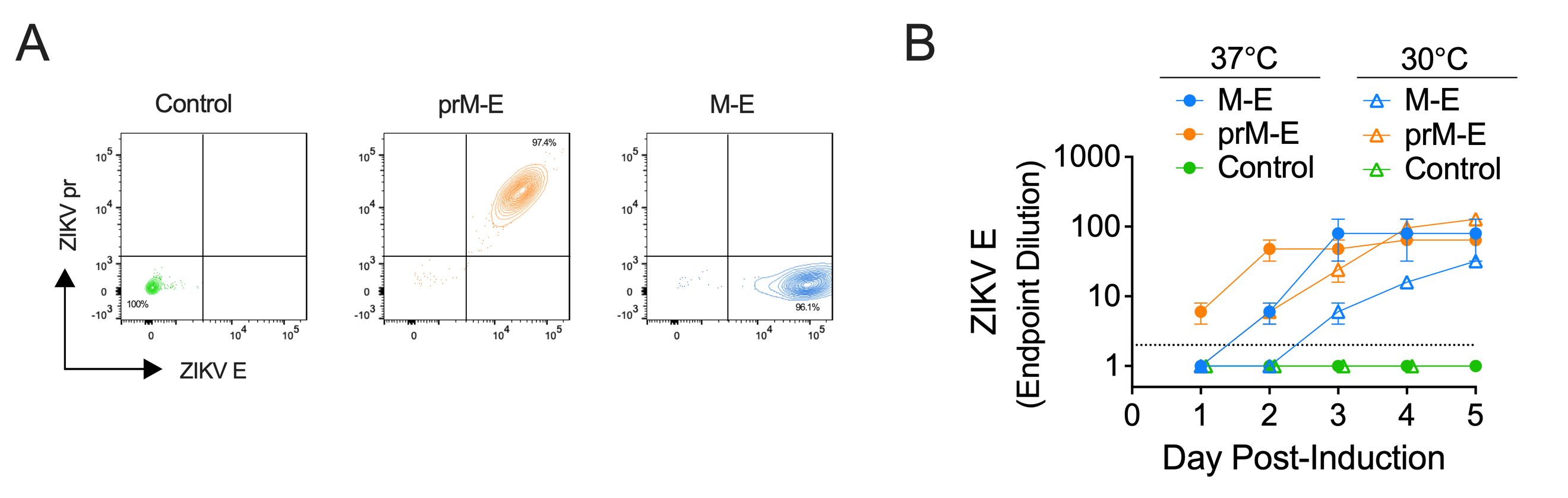

**Fig. S1. Inducible cell lines expressing ZIKV prM-E or M-E. (A)** ZIKV prM-E expressing, M-E expressing, or control CHO cells were subjected to intracellular staining 2 days post-induction with tetracyline. Cells were stained with a ZIKV pr-specific mAb (19pr) conjugated to Alexa Fluor-488 and a ZIKV E protein-specific mAb (ZV-67) conjugated to Alexa Fluor-647 to confirm expression. **(B)** ZIKV E protein ELISA endpoint dilutions for supernatants sampled from tetracycline-induced CHO cell lines incubated in parallel at 30°C and 37°C. Data is the average of two independent experiments. Error bars indicate the range. The dotted line indicates the starting dilution factor of 2; samples that were not positive at 1:2 were assigned an endpoint dilution of 1. The 37°C data for M-E and prM-E expressing cells is also shown in **Fig. 1D**.

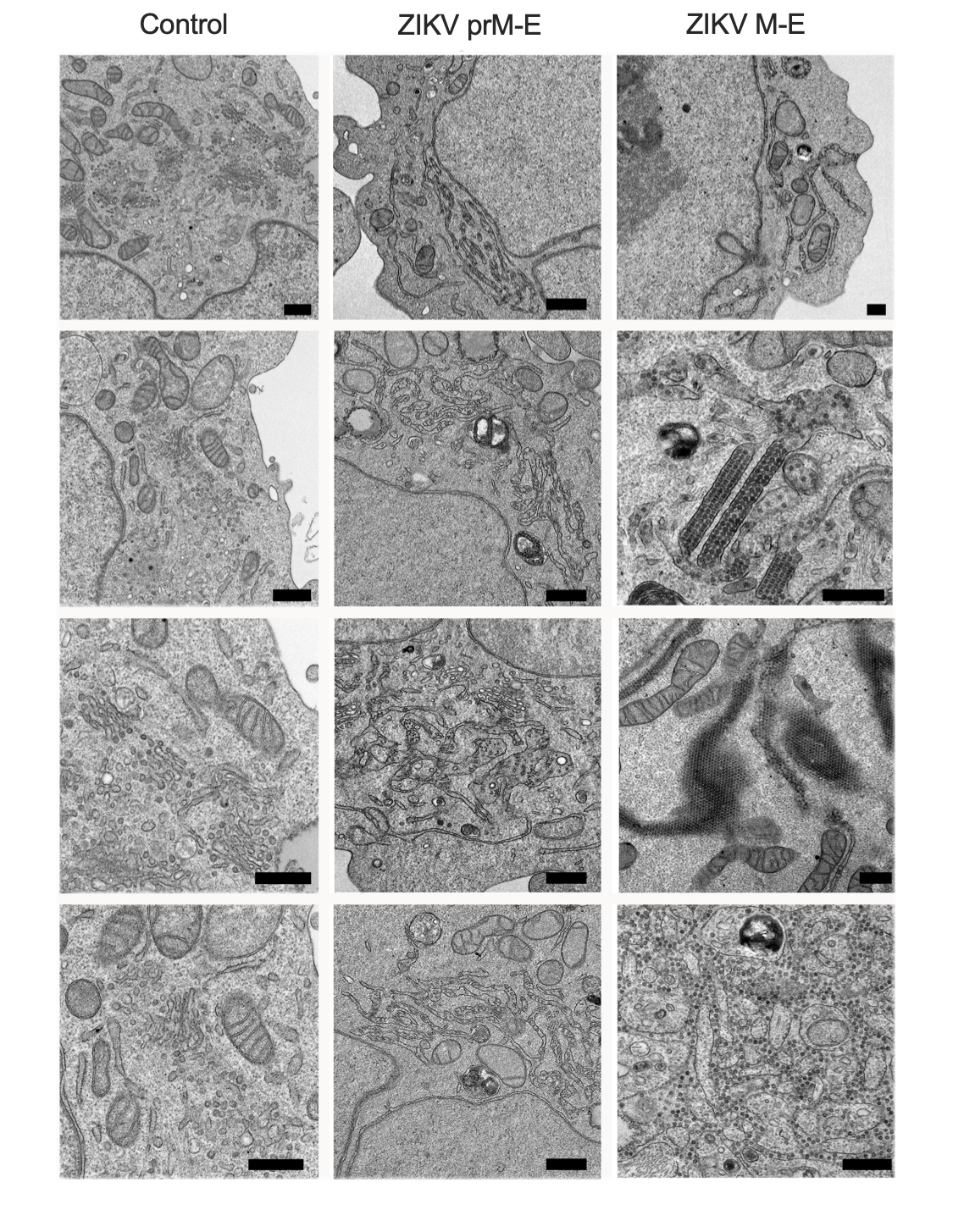

**Fig. S2**. **Additional images of ZIKV subviral particles in CHO cell lines engineered to express ZIKV prM-E and M-E.** Transmission electron microscopy images of cells 2 days post-induction with tetracyline. Scale bars represent 0.5 microns.

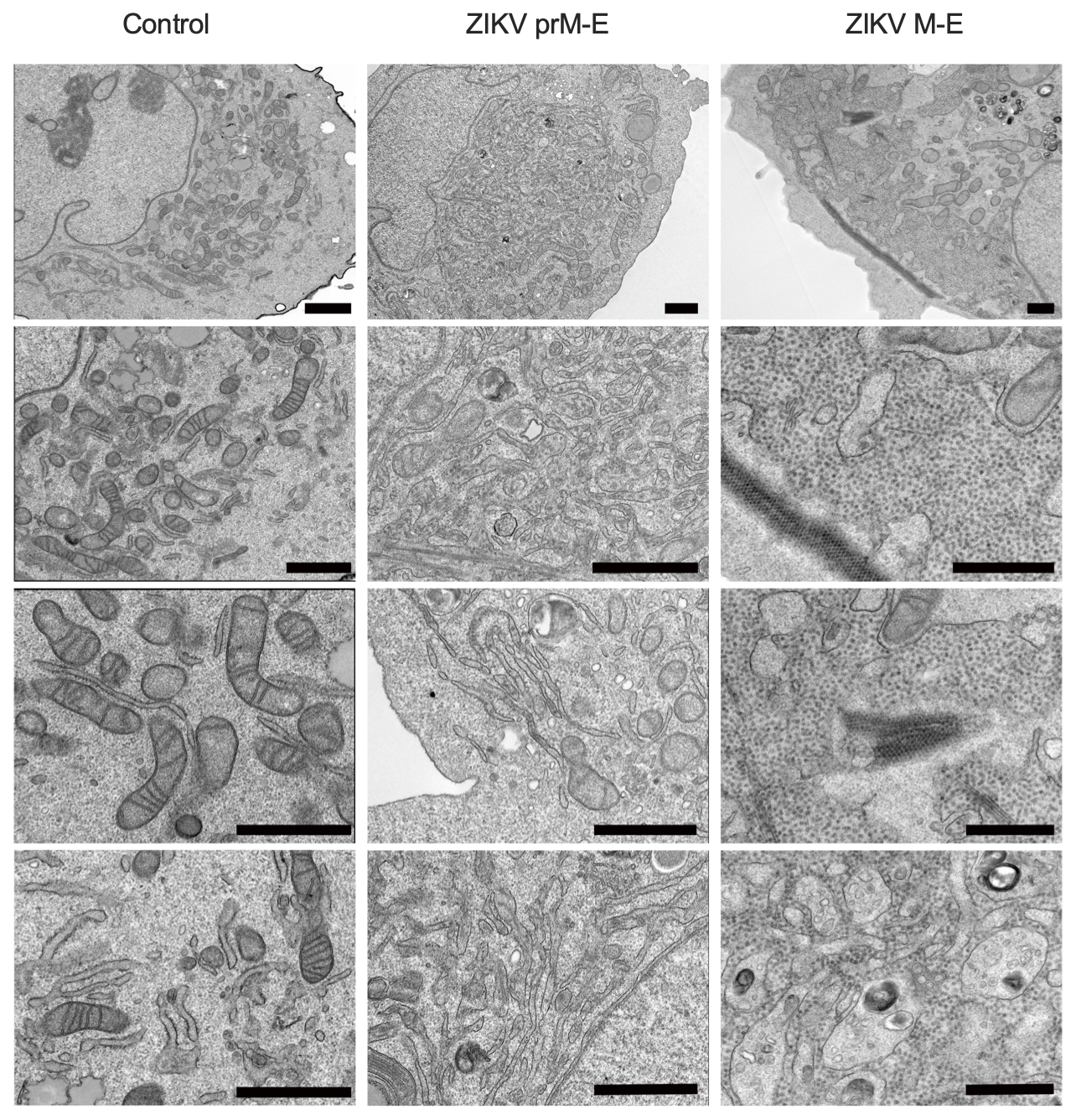

**Fig. S3**. **Images of ZIKV prM-E and M-E expressing CHO cell lines on day 4 post-induction.** Transmission electron microscopy images of cells 4 days post-induction with tetracycline. Scale bars represent 1 micron.

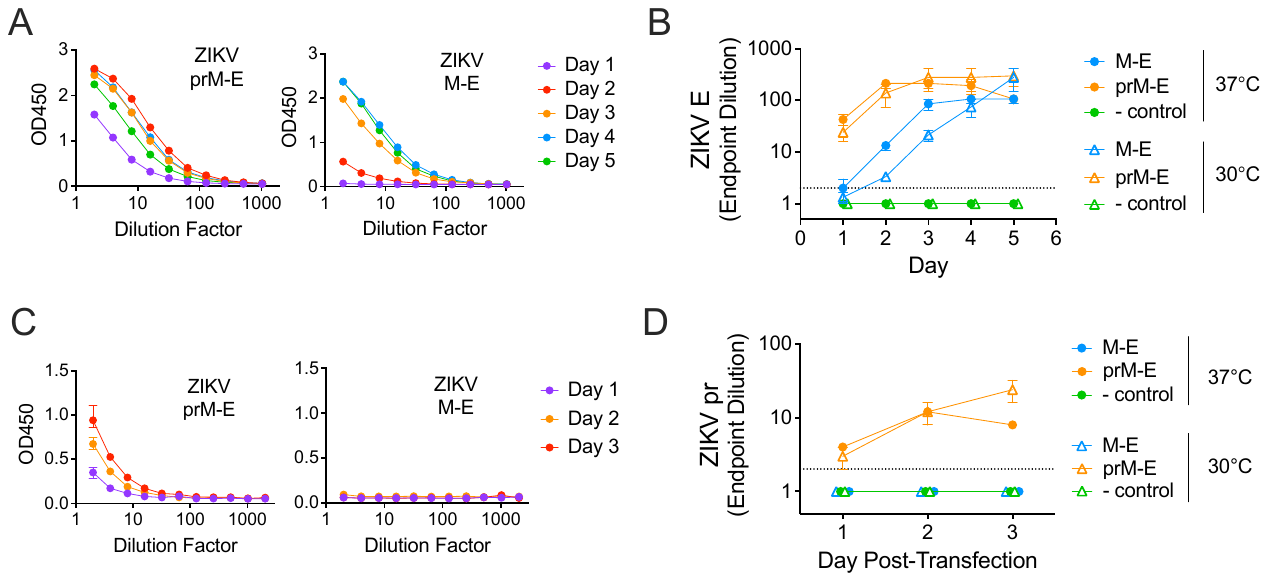

**Fig. S4. Secretion of ZIKV E and prM protein in transfected cells.** ZIKV E (**A-B**) and prM (**C-D**) protein secretion was measured by ELISA following transfection of 293T cells with plasmids encoding ZIKV prM-E, ZIKV M-E, or empty vector. Cells were incubated at 30°C or 37°C, and supernatant was sampled on the indicated days post-transfection. (**A**) Representative E protein ELISA dose-response curves from a 37°C experiment are shown, and the data used to calculate endpoint dilutions for each day. Error bars indicate the range of duplicate technical replicates. (**B**) ELISA endpoint dilutions for 30°C and 37°C transfection experiments performed in parallel. Data are the average of three independent experiments. Error bars indicate the standard error. (**C**) Representative pr protein ELISA dose-response curves from a 37°C experiment are shown, and the data used to calculate endpoint dilutions for each day. Error bars indicate the range of quadruplicate technical replicates. (**D**) ELISA endpoint dilutions for 30°C and 37°C transfection experiments performed in parallel. Data are the average of two independent experiments. Error bars indicate the range. The dotted line indicates the starting dilution factor of 2; samples not positive at a 1:2 dilution were assigned a value of 1.

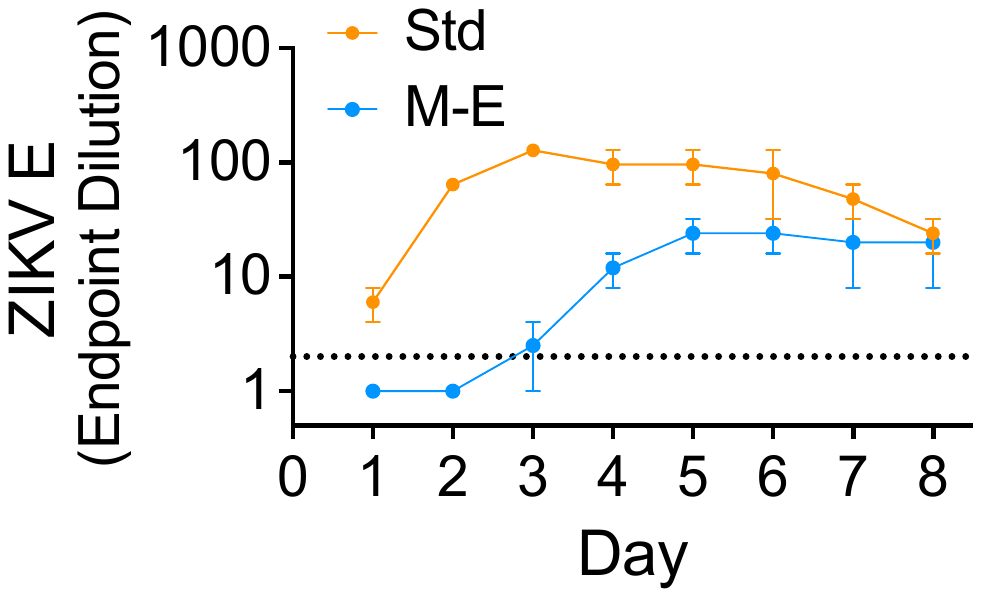

**Fig. S5**. **Secretion of ZIKV E protein from RVP-producing cells.** ZIKV RVPs capable of a single round of infection were generated by complementation of a GFP-expressing subgenomic replicon with a plasmid encoding ZIKV C, and a third plasmid encoding either ZIKV prM-E (Std) or M-E. Shown are ELISA endpoint dilutions for supernatants sampled from transfected 293T cells on days 1-8 post transfection. Data are the average of two independent experiments (see Fig. 2A for infectivity data from the same experiments). Error bars indicate the range. The dotted line indicates the starting dilution factor of 2; samples not positive at a 1:2 dilution were assigned an endpoint dilution of 1.

**Movie S1.** A tomogram of a 150 nm semi-thick section from ZIKV M-E expressing cells showing both individual and densely packed SVPs found in membrane-bound structures.
